# Reversed depth in anti-correlated random dot stereograms and central-peripheral difference in visual inference

**DOI:** 10.1101/225532

**Authors:** Li Zhaoping, Joelle Ackermann

## Abstract

Two images of random black and white dots, one for each eye, can represent object surfaces in a threedimensional scene when the dots correspond interocularly in a random dot stereogram (RDS). The spatial disparities between the corresponding dots represent depths of object surfaces. If the dots become anti-correlated such that a black dot in one monocular image corresponds to a white dot in the other, disparity-tuned neurons in the primary visual cortex (V1) respond as if their preferred disparities become non-preferred and vice versa, thereby reversing the disparity signs reported to higher visual areas. Humans have great difficulty perceiving the reversed depth, or any depth at all, in anti-correlated RDSs. We report that the reversed depth is more easily perceived when the RDSs are viewed in peripheral visual field, supporting a recently proposed central-peripheral dichotomy in mechanisms of feedback from higher to lower visual cortical areas for visual inference.

## 2 Introduction

Julesz (1971) first reported that one cannot see depth or reversed depth in anti-correlated RDSs, although one can see veridical depths when the density of dots in the anti-correlated RDSs is sufficiently sparse (Cumming et al., 1998) or in simple stereograms of reversed-contrast between the eyes(Helmholtz, 1925; Cogan et al., 1995). Fig. 1 shows three RDSs, each representing two depth planes, one for a central disk (of radius *r*) at disparity *d* ≠ 0 and the other contained the surrounding ring of dots at zero disparity and within radius *R* > *r*. The RDS in Fig. 1A is correlated, the two monocular images were identical to each other except for, (1), the dots for the central disk in the left eye image were horizontally displaced from their corresponding dots in the right eye image by a disparity d, and, (2), some monocular dots were near the border between the two depth planes due to surface occlusion. The disk appears in front or behind the surrounding ring of dots depending on whether *d* > 0 or *d* < 0. If the receptive field of a disparity-tuned V1 neuron covers only the dots for the central disk, the neuron would give high or low responses when the disparity d is its preferred or non-preferred disparity, respectively. However, if each dot within the disk and its corresponding dot in the other eye have opposite contrasts, as in the anti-correlated RDS in Fig. 1C, the V1 neuron would respond as if the preferred disparity becomes non-preferred and vice versa (Cumming and Parker, 1997; Ohzawa et al., 1990). In this RDS, human observers usually cannot see the veridical depth, nor can they see the reversed depth reported by the V1 neurons to higher visual areas. Fig. 1B, the zero-correlated stereogram, is like Fig. 1A or Fig. 1C, except that, within the disk, each dot and its counterpart in the other monocular image have the same or opposite contrasts with equal probability. Veridical depth can be easily seen in this RDS by human observers (Doi et al., 2011).

**Figure 1:**
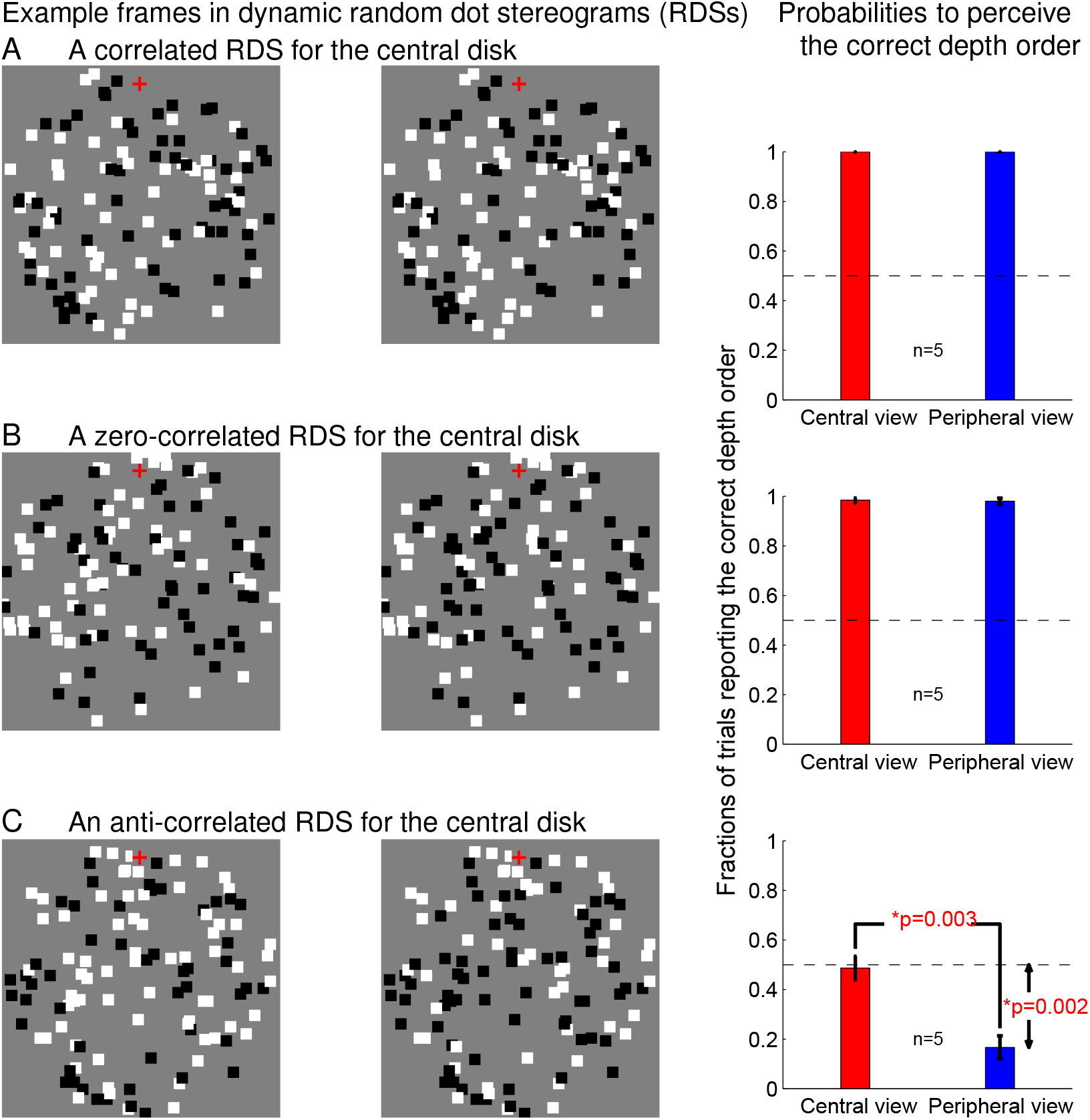
Examples of the RDSs (left) for two underlying depth planes and the corresponding observer performances (right) in reporting which depth plane was closer in one (Exp. 1) of the experiments. In each RDS (the pair of left and right monocular images), one depth plane of binocular disparity *d* = ±0.087° contains dots within a disk (radius *r* = 3°) at the center of the images, the other depth plane at zero disparity contains the surrounding dots (within radius *R* = 4.3°). All dots are squares with side length *s* = 0.348°. In the correlated (A), zero-correlated (B), or anti-correlated (C) RDS, the probability that each dot for the disk in one monocular image and its corresponding dot in the other monocular image have the same contrast (black or white) is *p* = 1, 0.5, or 0, respectively. For each surrounding dot outside the disk, this probability is always unity, not considering any monocular dots near the circumference of the disk due to local surface occlusions. The dashed line in the right column marks the probability (0.5) to correctly report the depth order between the two planes by chance. Observers fixated on the binocular (zero disparity) red cross (size *s_f_* = 0.44°) at *e*_o_ = 3.65° or *e*_1_ = 10.1° (not shown in the figure) above the center of the disk to view the RDSs centrally or peripherally.

However, previous studies on depth perception in anti-correlated stereograms focused on depth per ception in central visual field. Here we report that the reversed depth can be perceived in peripheral visual field. In our experiments, each frame in a dynamic RDS intended for the central viewing was like those in Fig. 1, adapted from those in a previous study(Doi et al., 2011). The black and white square dots had side length s. In each RDS, the dots would cover a fraction *f* = 0.25 of the image area for the disk and the ring if there were no dot occlusions. These RDSs were displayed dynamically for 1.5 second, each dichoptic image frame was refreshed every 100 milliseconds by an independent set of random dots given the stimulus parameters (i.e., *r, R, d, s, f*, and the correlation of the RDS). After this display period, observers took their time to report, by pressing one of the two assigned buttons in forced-choice, whether the disk of dots appeared in front or behind the surrounding ring of dots. When the RDSs were viewed in the central visual field, observers had to fixate on the binocular red cross of size *S_f_* at distance *e_o_* (with *r* < *e_o_* < *R*) above the center of the disk (Fig. 1); when the RDSs were viewed peripherally, observers fixating on a second binocular red cross added at a distance *e*_1_ > *R* above the disk center. Each experimental session randomly interleaved six types of trials: two fixation positions (for central and peripheral viewings), each with RDSs of three possible correlation levels (full-, zero-, and anti-correlation) as in Fig. 1. Only observers sufficiently able to correctly see the depth order in the fully-correlated RDSs (in practice trials) in both central and peripheral viewing were admitted to data taking. Part of our results was recently reported in abstract form (Zhaoping, 2017a).

## 3 Methods

The equipmental set up for this study copies that in a previous study (Zhaoping, 2017b). Each of the six conditions in an experimental session had 50 trials, in each trial the central disk was in front or behind the surrounding ring with equal probability. Each trial contained, in sequential order, a pre-fixation period which ended by observer’s button press, a fixation period, a period to display the RDSs, and finally a period for observer’s depth order report. The binocular red cross(es) displayed with the RDSs were present in each period. The cross among the ring of dots was at the same position on the display screen across the trials, whether it was the fixation cross (in centrally viewed trials) or not. During the pre-fixation period, observers were cued to the designated fixation cross by the positions of the two binocular text strings “press any button” and “for the next trial”, which were displayed to the left and right, respectively, of this cross until the start of the fixation period. Throughout each session, a black binocular rectangular frame was constantly on the display screen like in a previous study (Zhaoping, 2017b) to anchor vergence, enclosing the RDS region, the red crosses, and the texts. The display screen had the same gray background as the gray background in the RDSs. All the binocular items (the red crosses and the instruction texts) had the same binocular zero-disparity defined by the vergence anchoring frame. The fixation period was 0.7 second for observers who were not gaze tracked (this applied to one subject, the first author, who participated in all experiments, and to one session of another subject whose gaze was monitored by the experimenter through a video camera), otherwise, it ended at the first moment after 0.7 second into the fixation period when the eye tracker determined that the observer was fixating properly. All observers maintained their fixations properly in more than 90% of the trials. All observers other than one (the first author) were naive to the purpose of the study.

Before an experimental session, the observer went through the following four groups of practice trials sequentially: (1) 10 trials of the test condition with central viewing and fully correlated RDSs, in each trial the dynamic RDSs were modified to be a single static RDS viewed for two seconds to make the task easier; (2) 10 trials as in (1) but using the dynamic RDSs in the real experimental condition, (3) 20 trials, randomly interleaving 10 trials each of the two experimental conditions of fully correlated RDSs, one with central and one with peripheral viewing; (4) 6 example trials, one from each of the six experimental conditions. If the observer could correctly judge the depth order in no less than 90% of the trials in each condition of the practice trials in the groups (1), (2), and (3) above (roughly about 50% of observers could not achieve this, particular for the trials with peripheral viewing), he or she was then admitted to the experimental session. Since a previous study (Doi et al., 2011) indicated that observers could adequately perceive the depth order in zero-correlated RDSs (with central viewing), if an observer incorrectly reported the depth order in more than 25% of the experimental trials with zero-correlation RDSs by either fixation position, the data of this session was excluded from further analysis (one such session was therefore removed). Significant difference between the results for central and peripheral conditions were obtained by matched sample t-tests.

In Exps. 1–3, each RDS frame of the dynamic RDSs was made independently of any other frame as follows. Start with two concentric disks (radius *r* and *R* > *r*) of random dots. For each disk, each random dot was at any location within the disk with equal probability and was randomly black or white with equal probability; the total number of dots is such that the total image area covered by the dots would be 25% of the disk area if there were no mutual occlusions between the dots. The two monocular images of the RDS are made from these two disks of dots as follows. For the depth plane (the disk) of disparity *d*, the left and right monocular images contain the dots from the smaller disk (with radius *r*) after these dots were shifted horizontally by *d*/2 and –*d*/2, respectively. (The *d* is an integer multiple of image pixel size. When this integer is odd, the shift magnitude is (*d* + 1)/2 for one randomly chosen monocular image for each trial and (*d* – 1)/2 for the other monocular image.) For each monocular image, the dots for the surrounding ring were those from the larger disk (with radius *R*), excluding any dot that was either within the image area of the shifted smaller disk or would overlap in the monocular image with any dots from the smaller disk. The RDSs for Exp. 4/5 were made similarly, except that *R* = *R*_2_ and any dot in the larger disk was excluded if it was within radius *R*_1_ (*r* < *R*_1_ < *R*_2_) from the center of the disk or if it was monocular.

## 4 Results and Discussion

As shown in Fig. 1, with a disparity step of |*d*| = 0.087° between the two depth planes, observers could adequately judge the depth order when the dots for the central disk were full- or zero-correlated (*s* = 0.348°, *S_f_* = 0.44°, *r* = 3°, *R* = 4.3°). This was so both for when observers fixated centrally and peripherally, at, respectively, *e*_0_ – *r* = 0.65° and *e*_1_ – *r* = 7.1° above the closest depth discontinuity, in line with previous findings (Westheimer, 1982; Doi et al., 2011). However, with anti-correlated RDSs for the disk, observers’ performance was at chance by central view, as in previous studies (Cumming et al., 1998; Julesz, 1971) (also see Doi et al. (2011) for small d). However, by peripheral view, performance was significantly lower than chance, i.e., the observers were significantly better than chance to perceive the reversed depth order reported by V1 neurons (Qian, 1997), even though visual acuity is worse with peripheral viewing.

These results, partly shown again as Exp. 1 in Fig. 2, were from the first experiment in which central and peripheral viewing had the same visual input properties except for the retinal image locations for the dots and the additional red cross above and away from all the dots to serve as the fixation location for peripheral viewing. Relative to the visual acuity or the sizes of the receptive fields of the V1 cells responsive to the RDSs, the size of visual inputs for the RDSs was smaller for peripheral viewing. Scaling down the size of visual inputs by 50% for centrally viewed trials in two additional experiments, Exp. 2 and Exp. 3 in Fig. 2, did not impact the depth perception substantially. In Exp. 2, the scaling down was applied also to the disparity step size |*d*|; in Exp. 3, this |*d*| was instead unchanged and doubled (from the |*d*| in Exp. 1), respectively, for centrally and peripherally viewed trials. Hence, the visual input size is not critical for our results. Another two experiments, Exp. 4 and Exp. 5 in Fig. 2, showed that the central-peripheral difference was not due to the complexity of visual inputs at the boundary between the two depth surfaces, such as the monocular dots or, as suggested to the first author by Bruce Cumming, a mixture of correlated and anti-correlated dots that could fall within a single neural receptive field. Exp. 4 and Exp. 5 were like Exp. 1 and Exp. 2, respectively, however, a blank ring (of background luminance) in each image separated the disk of dots from the surrounding ring of dots and no monocular dots were present.

**Figure 2:**
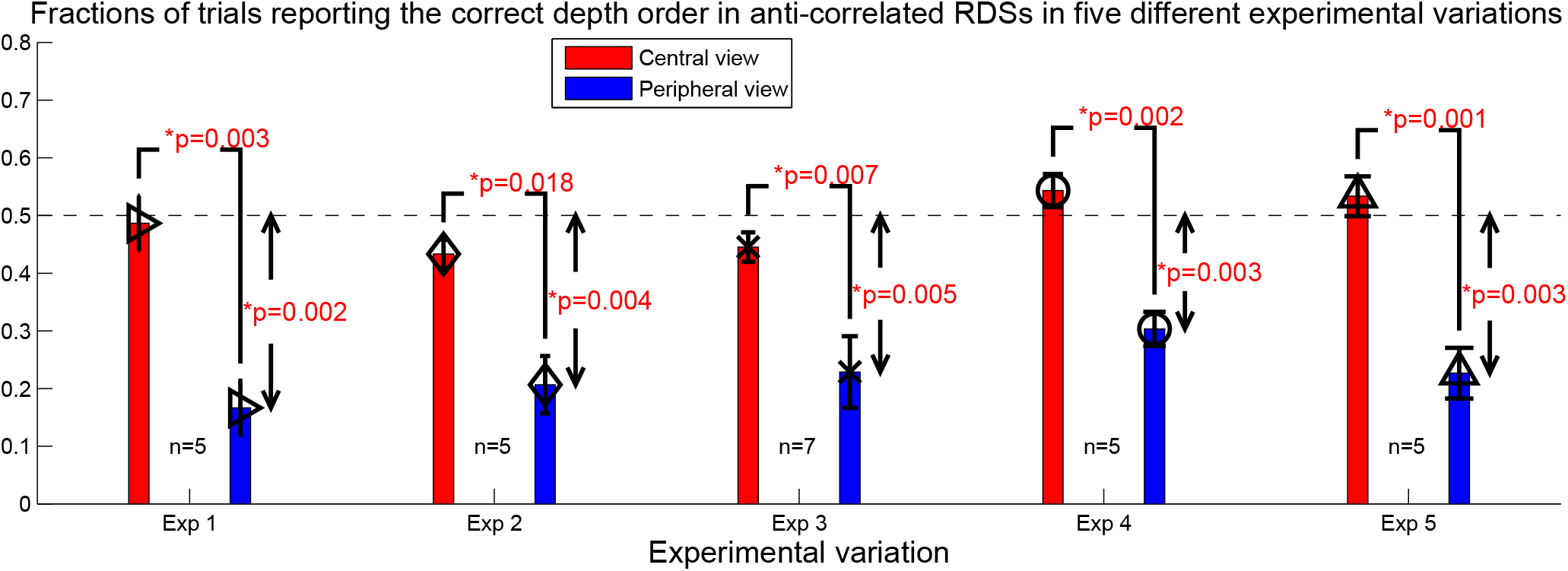
Tentency to perceive the reversed depth was stronger by peripheral than central viewing regardless of the sizes of the visual stimuli or the presence of a gap between the image regions for the two depth surfaces. Shown are observer performances for anti-correlated RDSs in five different experiments: Exps. 1-5, with data points ‘▻’, ‘◊’, ‘X’, ‘O’, and ‘Δ’, respectively. Exp 1 is the one whose results were in Fig. 1. Exp 2 differs from Exp. 1 by halving the spatial sizes of all the visual elements (*r, R, e_o_, s*, and *S_f_*) and disparity step (|*d*| = 0.0435^o^) for centrally viewed trials. Exp. 3 differs from Exp. 2 by doubling the disparity step in central/peripheral viewing, making |*d*| = 0.087^o^/0.17^o^. Exp. 4 differs from Exp. 1 by inserting a dot-free background ring of thickness 0.65^o^ in image space between the disk of dots and the surrounding ring of dots; accordingly, the inner and outer radiuses of the surrounding ring (of dots) were *R*_1_ = 3.7^o^ and *R*_2_ = 4.7^o^, the red cross in the surrounding ring was e_o_ = 4.1^o^ above the disk center, and no monocular dots were admitted. Exp. 5 differs from Exp. 4 by halving all the spatial scales (*r, R*_1_, *R*_2_, *e_o_*, *s*, *S_f_*, and |*d*|) of visual inputs for centrally viewed trials.

## 5 Discussion

Our study was designed to test a recent proposal (Zhaoping, 2017b) that perceptual inference employs a hierarchical feedback in central vision for the computation of analysis-by-synthesis and that this feedback is weaker or absent in peripheral vision. In this proposal, feedforward signals from V1 responses (e.g., to an input binocular disparity) suggest to higher visual areas an initial hypothesis for a perceptual decision (e.g., on the depth of an underlying object surface). If perception is unclear or ambiguous in central vision, the initial hypothesis is re-evaluated as follows. First, according to an internal model or prior knowledge of the visual world, the higher brain centers synthesize a would-be visual input that should resemble the actual visual input if the hypothesis is correct. Second, the synthesized input is fed back to lower visual areas such as V1 to compare with the actual visual input. Third, the strength of the initial perceptual hypothesis is reweighted according to the degree of the match between the synthesized and the actual inputs, such that a good or poor match strengthens or weakens, respectively, the initial hypothesis for the ultimate perceptual outcome. This inference process is called the feedforward-feedback-verify-weight (FFVW) process (Zhaoping, 2017b), and let us apply it to depth perception of our RDSs. When input contains an anti-correlated RDS of disparty d, V1 neurons respond as if the input disparity is –*d* (Cumming and Parker, 1997) and feed forward an initial hypothesis of a reversed disparity – *d*. Higher brain centers generate the synthesized inputs of disparity – *d*, and, according to the internal model of the visual world, the synthesized inputs to the left eye are correlated rather than anti-correlated with that to the right eye. The synthesized inputs are fed back to V1 and they do not match the anti-correlated actual inputs as represented by the population V1 responses. Consequently, the initial hypothesis of a reversed depth (by the – *d* disparity) fails the verification, is weakened or vetoed by the re-weighting, making it difficult for this reversed depth to be the ultimate perceptual outcome. The proposal states that feedback is weaker in the peripheral visual field, motivated by recent observations of tilt, motion, and color perception (Zhaoping, 2017b). Accordingly, the initial hypothesis of a reversed depth in the peripheral visual field is not easily verified or vetoed, and is thus more likely to become the perceptual outcome.

The central-peripheral dichotomy by the FFVW process should apply to general cases of visual inference about properties of visual objects. The previous work (Zhaoping, 2017b) used ambiguous perceptions of motion direction, tilt, and color of gratings to infer or reveal this dichotomy, and the current study provides a supporting evidence for this dichotomy. We can relate the reversed depth perception in anti-correlated RDSs with its motion analog, the reverved phi motion, when the perceived motion direction is opposite to that of the physical displacement between a picture and its photo-negative when the former is followed by the (slightly displaced) latter via a temporal dissolve (Anstis, 1970; Anstis and Rogers, 1975). Using the analog of the temporal dissolve in stereo vision, one can see the reversed depth in a stereogram if one eye views a picture and the other eye views a weighted average of this original image and its slightly displaced photo-negative (Rogers and Anstis, 1975). Just as reversed depth is difficult to see in central vision in anticorrelated stereograms, for reversed phi stimuli, the perceived motion direction is indeterminate by central viewing when the picture and its (displaced) photo-negative were presented successively without a temporal dissolve (Anstis and Rogers, 1975). It has also been noted that reversed phi motion is more easily seen in peripheral visual field (Anstis, 1970; Anstis and Rogers, 1975), while the physical motion direction is more easily seen in central visual field (Lu and Sperling, 1999), manifesting the central-peripheral dichotomy. Future studies are needed to explore other perceptual varieties in which the central-peripheral dichotomy in visual inference can apply, and to reveal neural mechanistic details of the FFVW process.

## 6 Acknowledgement

This study is supported by the Gatsby Charitable Foundation.

